# RELA is Sufficient to Mediate Interleukin-1 (IL-1) Repression of Androgen Receptor Expression and Activity in LNCaP Disease Progression Model

**DOI:** 10.1101/775320

**Authors:** Shayna E. Thomas-Jardin, Haley Dahl, Mohammed S. Kanchwala, Freedom Ha, Joan Jacob, Reshma Soundharrajan, Monica Bautista, Afshan F. Nawas, Dexter Robichaux, Ragini Mistry, Vanessa Anunobi, Chao Xing, Nikki A. Delk

## Abstract

**Background:** The Androgen Receptor (AR) nuclear transcription factor is a therapeutic target for prostate cancer (PCa). Unfortunately, patients can develop resistance to AR-targeted therapies and progress to lethal disease, underscoring the importance of understanding the molecular mechanisms that underlie treatment resistance. Inflammation is implicated in PCa initiation and progression and we have previously reported that the inflammatory cytokine, interleukin-1 (IL-1), represses *AR* mRNA levels and activity in AR-positive (AR^+^) PCa cell lines concomitant with the upregulation of pro-survival biomolecules. Thus, we contend that IL-1 can select for AR-independent, treatment-resistant PCa cells.

**Methods:** To begin to explore how IL-1 signaling leads to the repression of *AR* mRNA levels, we performed comprehensive pathway analysis on our RNA sequencing data from IL-1-treated LNCaP PCa cells. Our pathway analysis predicted Nuclear Factor Kappa B (NF-κB) p65 subunit (RELA), a canonical IL-1 signal transducer, to be significantly active and potentially regulate many genes, including *AR*. We used siRNA to silence the NF-κB family of transcription factor subunits, *RELA*, *RELB*, *c-REL*, *NFKB1*, or *NFKB2*, in IL-1-treated LNCaP, C4-2, and C4-2B PCa cell lines. C4-2 and C4-2B cell lines are castration-resistant LNCaP sublines and represent progression towards metastatic PCa disease; and we have previously shown that IL-1 represses *AR* mRNA levels in C4-2 and C4-2B cells.

**Results:** siRNA revealed that RELA alone is sufficient to mediate IL-1 repression of *AR* mRNA and AR activity. Intriguingly, while LNCaP cells are more sensitive to IL-1-mediated repression of *AR* than C4-2 and C4-2B cells, *RELA* siRNA led to a more striking de-repression of *AR* mRNA levels and AR activity in C4-2 and C4-2B cells than in LNCaP cells.

**Conclusions:** These data indicate that there are RELA-independent mechanisms that regulate IL-1-mediated *AR* repression in LNCaP cells and suggest that the switch to RELA-dependent IL-1 repression of *AR* in C4-2 and C4-2B cells reflects changes in epigenetic and transcriptional programs that evolve during PCa disease progression.

## INTRODUCTION

Androgen deprivation is used as a first line therapy for PCa patients, but some patients will inevitably progress to androgen-independent PCa^1^. Activating mutations, splice variation, or overexpression of AR can render AR ligand-independent or functional under castrate levels of androgen^1^. Therefore, patients that develop androgen independence are given anti-androgens that directly bind and prevent AR transactivation activity^1^. PCa tumors can, however, evolve anti-androgen resistance, leading to lethal disease^1^. Anti-androgens can function as AR agonists resulting in treatment resistance^1^; but anti-androgen resistance and AR independence can also result from the emergence of PCa tumor cells with low or no AR (AR^low/-^) accumulation or activity^2^.

We and others have found that IL-1 represses AR levels and activity in PCa cells^3–7^ and one group has shown that chronic IL-1 exposure reduces sensitivity to the anti-androgen, bicalutamide, in the androgen-dependent LNCaP PCa cell line^5^. While inflammation and IL-1 are associated with reduced AR levels or activity in benign prostatic hyperplasia^7^ and there is evidence that IL-1 inversely correlates with AR activity in metastatic PCa patient tissue^8^, it is unknown if IL-1 is sufficient to drive AR independence or treatment resistance during PCa disease progression. We previously reported that concomitant with AR repression, IL-1 induces a suite of genes in LNCaP cells that mimic basal gene expression in AR-independent PCa cell lines^6^, suggesting that IL-1 exposure could select for viable AR-independent PCa tumor cells. For example, we demonstrated that Sequestome-1 (p62), an autophagy and antioxidant protein from our gene suite^6^, is required for cell survival in AR-independent PCa cell lines^9^ and others have recently discovered that *p62* overexpression can drive prostate cell tumor formation *in vivo*^10^.

To begin to understand how IL-1 drives AR independence, we evaluated IL-1 regulation of *AR* repression across a disease progression cell line model using LNCaP, C4-2, and C4-2B PCa cell lines. LNCaP is an androgen-dependent cell line originating from a PCa patient lymph node metastasis. LNCaP cells require androgen and AR activity for survival and do not form metastatic tumors in mice^11–13^. The C4-2 and C4-2B PCa cell lines are LNCaP sublines isolated from a subcutaneous tumor (C4-2) in a castrated mouse that metastasized to bone (C4-2B) when re-grafted in a castrated mouse^13^. Thus, C4-2 and C4-2B cells represent progressive, metastatic, androgen-independent PCa disease^13^. As with LNCaP cells, IL-1 represses AR levels in C4-2 and C4-2B cell lines^6, 14^ and, thus, are relevant cell line models to begin to address IL-1 regulation of AR during disease progression.

Nuclear Factor Kappa B (NF-κB) transcription factor is the canonical mediator of IL-1 signaling^15^. The NF-κB transcription factor family is comprised of RELA (p65), RELB, c-REL, NFKB1 (p105/p50) and NFKB2 (p100/p52)^15^. RELA, RELB, and c-REL contain DNA binding and transactivation domains^15^. NFKB1 and NFKB2 are processed from 105 and 100 kDa proteins to 50 and 52 kDa subunits, respectively, that bind DNA but lack transactivation domains^15^. The NF-κB family members form homo- and heterodimers to induce or repress transcription^15^, where the RELA/p50 heterodimer typically regulates gene expression in response to IL-1^15^. Using RNA sequencing pathway analysis of IL-1-treated LNCaP cells, we identified several candidate transcriptional regulators of *AR* expression. Not surprisingly, our analysis identified RELA as one of the most significantly activated transcriptional regulators of *AR* expression in IL-1-treated cells predicted by IPA (z-score = 5.58, p-value 1.6E-11) and we show that RELA can indeed mediate IL-1 repression of *AR* levels and AR activity. Using the LNCaP PCa cell line progression model, our results also suggest that RELA-dependent repression of *AR* evolves with PCa progression towards androgen independence.

## MATERIALS AND METHODS

### Cell Culture

LNCaP, C4-2, and C4-2B prostate cancer (PCa) cell lines were maintained in a 37°C, 5.0% (v/v) CO2 growth chamber and cultured in DMEM (Gibco/Thermo Scientific; 1185-092) supplemented with 10% (v/v) fetal bovine essence (fetal bovine serum alternative) (FBE; Seradigm; 3100-500), 0.4mM L-glutamine (L-glut; Gibco/Invitrogen; 25030-081), and 10U/ml penicillin G sodium and 10 mg/ml streptomycin sulfate (pen-strep; Gibco/ Invitrogen; 15140-122).

### Cell Treatments

*<u>Intereleukin-1 (IL-1):</u>* Vehicle control (0.1% bovine serum albumin (BSA) in 1X phosphate buffered saline (PBS)), 0.5 ng/ml, 5 ng/ml, or 25 ng/ml IL-1α (Gold Bio, 1110-01A-10) or IL-1β (Gold Bio, 1110-01B-10) were added to DMEM/10% FBE growth medium. *<u>Gene silencing (siRNA):</u>* The following siRNA concentrations were used: 70nM non-targeting siRNA (Dharmacon, D-001206-13-05), *RELA* siRNA (Dharmacon, M-003533-02-0005), *NFKB1* siRNA (Dharmacon, M-003520-01-0005) or *p62* siRNA (Dharmacon, M-010230-00-0005); 100nM non-targeting siRNA, *RELB* siRNA (Dharmacon, M-004767-02-0005), *c-REL* siRNA (Dharmacon, M-004768-01-0005), or *NFKB2* siRNA (Dharmacon, M-003918-02-0005); 70nM non-targeting siRNA (Origene, SR30004) or *RELA* siRNA (Origene, SR 321602A-C). Cells were transfected with siRNA using siTran 1.0 transfection reagent (Origene, TT300003) added to cells in DMEM/10% FBE growth medium for 1 day. siRNA containing medium was then removed and replaced with fresh DMEM/10% FBE plus vehicle control, IL-1α, or IL-1β. *<u>Serum starvation:</u>* After plating, cells were maintained in DMEM/2.5% FBE for several days prior to replacing medium with DMEM/0% FBE (serum starvation) or DMEM/10% FBE (replete medium control).

### Cell Counts

Cells were fixed in 100% cold methanol. The nuclei were stained with DAPI (Roche Diagnostics, 10236276001) and the stained cells imaged and counted using the Cytation3 Cell Imaging Multi-Mode Reader (BioTek, Winooski, Vermont).

### Crystal Violet Staining

Cells were fixed in 100% cold methanol at −20C for 30 minutes and stained with 0.25% crystal violet (Acros Organics, AC212120250) in 10% methanol for 5 minutes at room temperature. Excess stain was washed off with deionized water.

### RNA Isolation and Reverse Transcription Quantitative PCR (RT-QPCR)

Total RNA was extracted, reverse transcribed, and analyzed by RT-QPCR as previously described^6^. Primer sequences for genes of interest are listed below. Gene of interest cycle times (CT) were normalized to the *β-actin*. Relative mRNA levels were calculated using the 2-ΔΔCT method. Primer sequences, 5’-3’: *Androgen Receptor (AR)*, forward AAGACGCTTCTACCAGCTCACCAA, reverse TCCCAGAAAGGATCTTGGGCACTT; *Beta actin (β-actin)*, forward GATGAGATTGGCATGGCTTT, reverse CACCTTCACCGGTCCAGTTT; *Prostate Specific Antigen (PSA)*, forward CACCTGCTCGGGTGATTCTG, reverse ACTGCCCCATGACGTGATAC; *Sequestosome-1 (p62)*, forward AAATGGGTCCACCAGGAAACTGGA, reverse TCAACTTCAATGCCCAGAGGGCTA; *Mitochondrial Superoxide Dismutase 2 (SOD2)*, forward GGCCTACGTGAACAACCTGA, reverse GTTCTCCACCACCGTTAGGG; *RELA*, forward TGAACCAGGGCATACCTGTG, reverse CCCCTGTCACTAGGCGAGTT; *NFKB1*, forward TGATCCATATTTGGGAAGGCCTGA, reverse GTATGGGCCATCTGTTGGCAG; *NFKB2*, forward CCTAAGCAGAGAGGCTTCCG, reverse TCTTTCGGCCCTTCTCACTG; *RELB*, forward AGCTCTACTTGCTCTGCGAC, reverse AACACAATGGCAATCTGGCG; *c-REL*, forward ACAGCACAGACAACAACCGA, reverse AGTAGCCGTCTCTGCAGTCT.

### RNA Sequencing (RNA-seq) Analysis

Pathway analysis was conducted using QIAGEN’s Ingenuity Pathway Analysis (IPA) tool (http://www.qiagen.com/ingenuity). RNA-seq datasets used for this study are available at GEO NCBI, accession GSE105088. LNCaP samples treated with vehicle control or IL-1β were considered and 3,658 differentially expressed genes with log2 counts per million (CPM) ≥ 0, absolute log2 fold change (FC) > 0.6, and false discovery rate (FDR) ≤ 0.05 were considered for further downstream analysis.

### Western Blot

Western blotting was done as previously described^6^. Briefly, primary and secondary antibodies were diluted in 2.5% BSA in 1X TBST. Protein blot bands were visualized using SuperSignal West Femto Maximum Sensitivity Substrate (Thermo Scientific; 34095) or Clarity Western ECL Substrate (Bio-Rad; 1705061) and imaged using the Amersham Imager 600 (GE, Marlborough, MA). *<u>Primary antibodies:</u>* AR (Cell Signaling; D6F11), p62 (Abnova; H00008878-M01), SOD2 (Abgent; AM7579a), β-actin (Santa Cruz; sc-69879), RELA (Cell Signaling; 6956S), NFKB1 (Cell Signaling; 3035S), PSA (Cell Signaling; 5365S), or c-REL (Cell Signaling; 4727T). *<u>Secondary antibodies:</u>* Sheep anti-mouse (Jackson ImmunoResearch Laboratories; 515-035-062), goat anti-rabbit (Abnova; PAB10822). *<u>Densitometry:</u>* Western blot densitometry for protein bands was performed using Image J (National Institutes of Health, Bethesda, Maryland). Protein bands were first normalized to β-actin loading control and then normalized to the treatment control.

### Immunofluorescence

Cells were fixed and permeabilized with 100% methanol at −20°C for 30 minutes. Fixed cells were blocked with 2.5% BSA in 1X PBS at room temperature for at least 30 minutes. Antibodies were diluted in 2.5% BSA in 1X PBS. *<u>Primary antibody:</u>* AR (Cell Signaling; D6F11). *<u>Secondary antibody</u>*: Alexafluor 568, goat anti-rabbit (Invitrogen; A11061). Nuclei were stained with DAPI (Roche Diagnostics, 10236276001). Immunostained cells were imaged at 20X magnification using the Nikon Eclipse Ti fluorescence microscope and Nikon NIS Elements software (Nikon, Melville, New York).

### Statistical Analysis

Statistical significance was determined using unpaired student’s t-test. p-value ≤ 0.05 is considered statistically significant. Error bars, +/- standard deviation (STDEV); n = 3 biological replicates. Experiments were repeated at least 3 times and representative data shown.

## RESULTS

### RNA Sequencing pathway analysis reveals RELA as a potential regulator of *AR* expression

We previously performed RNA sequencing on the LNCaP PCa cell line treated with IL-1β and found that *AR* and AR target genes are down regulated^6^. Here, we performed upstream regulator analysis using IPA on the vehicle control versus IL-1β-treated LNCaP data sets and identified RELA as a significantly active transcriptional regulator of *AR* (z-score = 5.58, p-value 1.6E-11). Figure 1 shows selected RELA interactors from our IPA analysis. A complete list of RELA interactors identified based on IPA for IL-1β-treated LNCaP cells and their relationships can be found in Supplemental Table 1. It is worth noting that IPA’s knowledge base indicates RELA to be a positive regulator of *AR* expression which is inconsistent (indicated by the yellow arrow between RELA and AR, Fig. 1) with our RNA sequencing data that indicates that IL-1β represses *AR* mRNA levels in LNCaP cells. Therefore, we set out to experimentally test the relationship between RELA activity and *AR* expression in our model. To do so, we silenced *RELA* in IL-1-treated LNCaP cells and used *Super Oxide Dismutase 2* (*SOD2*) expression as a positive control for RELA activity.

**Figure 1.**
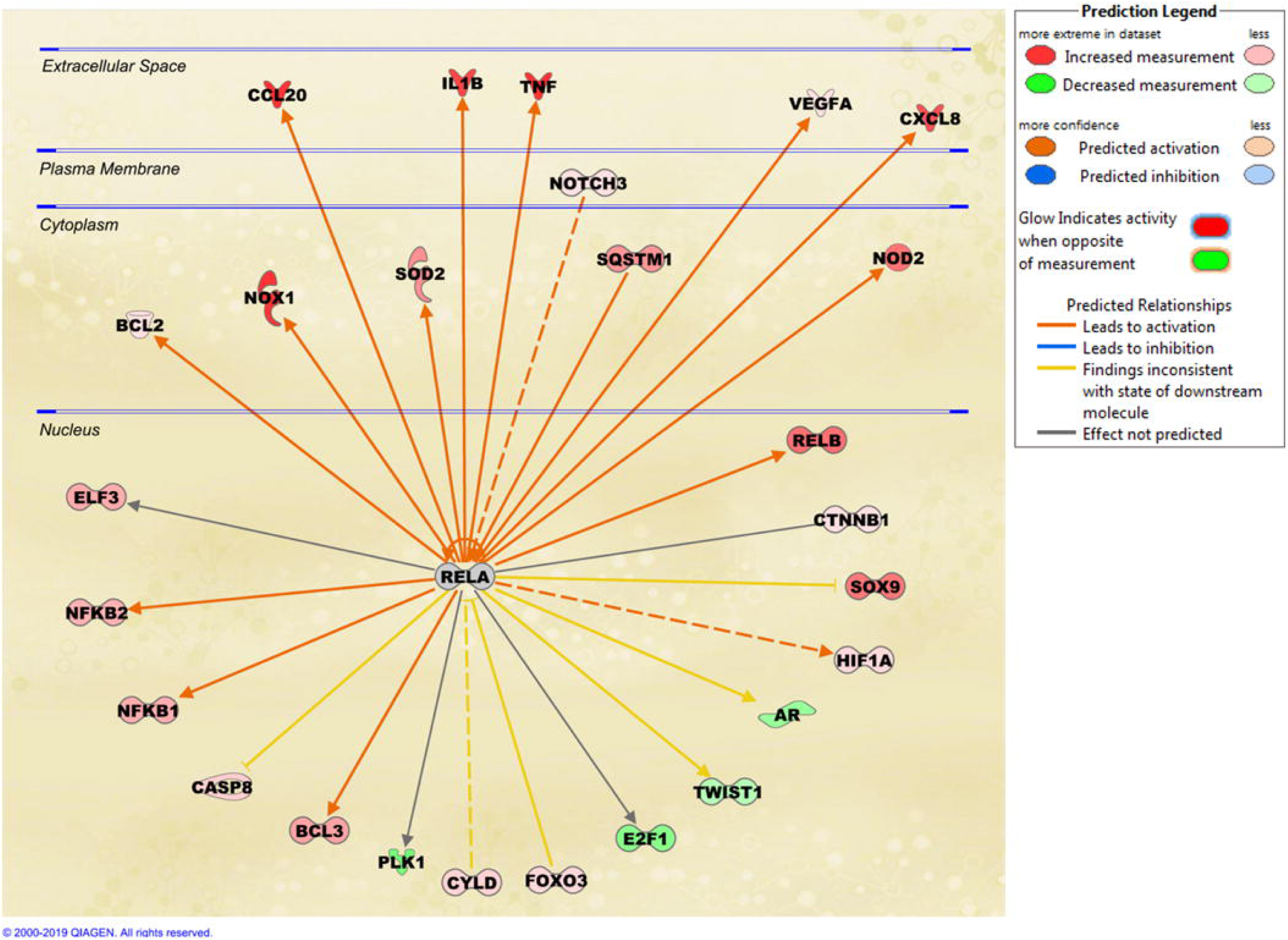
RELA Interactome. Upstream Regulator Analysis of IPA identified RELA as one of the most activated transcription factors (activation z-score = 5.58). Here we show selected interactome of RELA which has significant overlap (p-value 1.6E-11) with our gene set (GSE105088) of IL-1-induced genes in LNCaP. A complete list of proteins in this RELA interactome can be found in Supplemental Table 1. Red color indicates the gene encoding the protein was significantly upregulated (IL-1-induced) in our gene set and green color indicates downregulation (IL-1-inhibited). Orange color indicates predicted activation consistent with knowledgebase and blue color indicates predicted inhibition, yellow color indicated inconsistent findings between our data and the knowledgebase. Solid arrow indicates direct relationship and dotted line indicates indirect relationship. IPA knowledgebase predicts that RELA activates *AR* expression in IL-1-treated cells which is inconsistent (yellow arrow) with our gene expression data (GSE105088).

### LNCaP, C4-2 and C4-2B cell lines show differential sensitivity to IL-1

Prior to embarking on our investigation of RELA regulation of *AR* expression in the LNCaP cells, we had observed that LNCaP cells are more sensitive to IL-1 signaling than C4-2 or C4-2B cells. For example, we find that IL-1-treated LNCaP cells show greater SOD2 upregulation and AR repression than C4-2 or C4-2B cells. Based on RT-QPCR, 25 ng/ml IL-1α or IL-1β induced *SOD2* mRNA levels ∼18 fold in LNCaP cells versus ∼6 fold in C4-2B cells (Fig. 2A) and based on western blot, 25 ng/ml IL-1α or IL-1β induced SOD2 protein accumulation ∼5 fold in LNCaP cells versus ∼3 fold in C4-2 and C4-2B cells (Fig. 2B). Furthermore, 25 ng/ml IL-1α or IL-1β repressed *AR* mRNA levels ∼5 fold in LNCaP cells versus ∼3 fold in C4-2 or ∼2 fold in C4-2B cells (Fig. 2A). In kind, western blot shows that 25 ng/ml IL-1α or IL-1β repressed AR protein accumulation ∼6 fold in LNCaP cells versus ∼4 fold in C4-2 cells or ∼2 fold in C4-2B cells (Fig. 2B). Interestingly, while IL-1 receptor (IL-1R1) mRNA levels are elevated in C4-2 and C4-2B cells and RELA mRNA and protein levels are similar across LNCaP, C4-2, and C4-2B cell lines, basal *NFKB1* mRNA and p50 protein levels are lower in C4-2 and C4-2B cells than in LNCaP cells and IL-1-mediated induction of *NFKB1* mRNA and p50 protein levels is not as robust in C4-2 or C4-2B cells as in LNCaP cells (Fig. 2A & B). Given that canonical IL-1 signaling is mediated via the RELA/p50 heterodimer^15^ the lower *NFKB1* mRNA and p50 protein levels might underlie C4-2 and C4-2B reduced sensitivity to IL-1.

**Figure 2.**
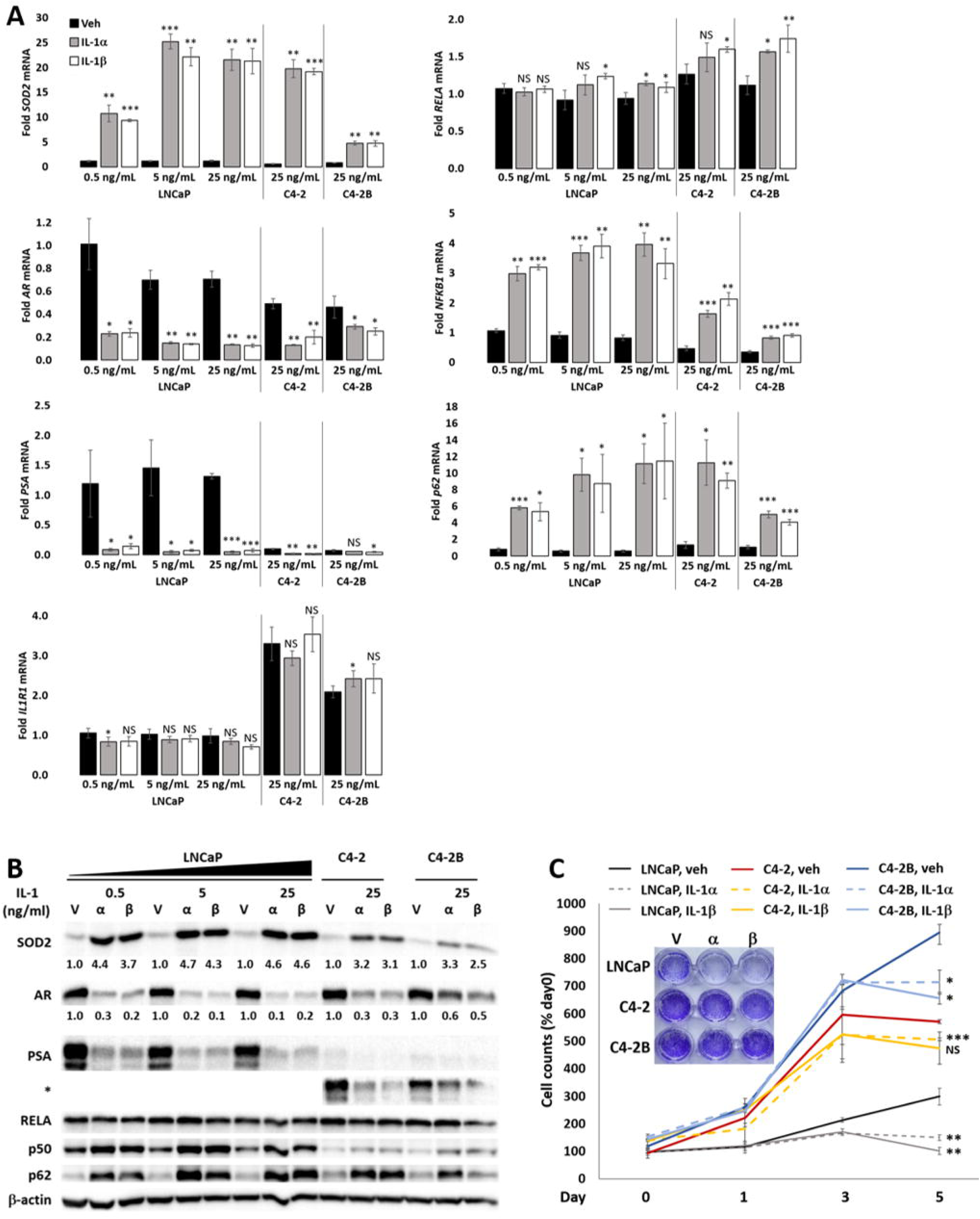
The LNCaP series shows differential sensitivity to IL-1. (A) RT-QPCR and (B) western blot analysis and densitometry were performed for LNCaP, C4-2, and C4-2B cells treated for 2 days with vehicle control (Veh, V), IL-1α, or IL-1β. IL-1 upregulates SOD2, NFKB1(p50), and p62 levels and represses AR and PSA levels. RELA levels do not change. *IL-1R1* mRNA levels are basal high, but NFKB1(p50) levels are basally low in C4-2 and C4-2B. (C) LNCaP, C4-2, and C4-2B cells were treated with vehicle control (V), 25 ng/ml IL-1α, or 25 ng/ml IL-1β for 0-5 days. Cells were stained with DAPI for cells counts or stained with crystal violet (inset image). LNCaP cells are more sensitive to IL-1 gene and protein regulation and cytotoxicity than C4-2 or C4-2B cells. Error bars, ± STDEV of 3 biological replicates; p-value, *≤0.05, **≤0.005, ***≤0.0005. mRNA fold change is normalized to the vehicle control for LNCaP 0.5 ng/ml IL-1 treatment for all cell lines and treatments. Cell counts were normalized to day 0, vehicle control. Day 0 is the day prior to treatment.

In addition to reduced sensitivity to IL-1 modulation of target mRNA and protein levels, C4-2 and C4-2B cells are less sensitive to IL-1 induced cytotoxicity. LNCaP, C4-2, and C4-2B cells were grown in 25 ng/ml IL-1α or IL-1β for 1,3, or 5 days and the cell number was recorded (Fig. 2C). Cells were also visualized on day 5 using crystal violet stain (Fig. 2C). C4-2 and C4-2B cells show faster growth rate than LNCaP cells in both vehicle and IL-1 treatment conditions; but when comparing cytotoxicity in IL-1, C4-2 and C4-2B cells show ∼14-23% reduction in cell numbers on day 5, while LNCaP cells show ∼ 58% reduction (Fig. 2C). Thus, compared to LNCaP cells, C4-2 and C4-2B cells have reduced sensitivity to IL-1.

As stated earlier, C4-2 and C4-2B cell lines are less dependent on androgen than LNCaP cells^13^. For example, compared to LNCaP cells, C4-2 and C4-2B cells have reduced sensitivity to growth arrest under serum-free growth conditions (Supplemental Fig. 1). Interestingly, relative to LNCaP cells, C-42 and C4-2B cells have reduced basal mRNA and protein levels of the AR target gene, *Prostate Specific Antigen* (*PSA*) (Fig. 2); but, have higher basal levels of the AR target gene, *Kallikrein* (*KLK2*) (Supplemental Fig. 3A). Indicating that the regulation of AR target genes is altered in C4-2 and C4-2B cells. Taken together, cell response to IL-1 signaling may evolve with androgen, including IL-1-RELA regulation of AR levels and activity, which we investigated below.

### RELA mediates IL-1 repression of AR levels and activity differentially in LNCaP versus C4-2 and C4-2B cells

To determine if RELA mediates IL-1 repression of AR, we siRNA-silenced *RELA* in IL-1-treated LNCaP, C4-2, or C4-2B cells and analyzed AR mRNA levels and protein accumulation. We also analyzed PSA or KLK2 mRNA and/or protein levels as surrogates for AR transcriptional activity and we immunostained cells for AR nuclear accumulation. LNCaP, C4-2, and C4-2B cells were treated with vehicle control, 25 ng/mL IL-1α, or 25 ng/mL IL-1β in the presence of non-targeting control siRNA or two different pools of *RELA* siRNA. *RELA* siRNA was sufficient to attenuate IL-1 repression of AR, PSA, and KLK2 mRNA and/or protein (Fig. 3, Supplemental Fig. 2 & 3B) and AR nuclear accumulation (Fig. 4) in C4-2 and C4-2B cells treated with 25 ng/ml IL-1. However, *RELA* siRNA was not sufficient to attenuate IL-1 repression of AR or showed only a slight effect on PSA or KLK mRNA and/or protein in LNCaP cells treated with 25 ng/ml IL-1 (data not show). Given the enhanced sensitivity of LNCaP cells to IL-1, we instead silenced *RELA* in LNCaP cells treated with a lower IL-1 concentration (0.5 ng/mL) and found that *RELA* siRNA was sufficient to attenuate IL-1 repression of AR, PSA, and KLK2 mRNA and/or protein levels; however, the LNCaP response to *RELA* siRNA was not consistent or as robust as for the C4-2 or C4-2B cell lines (Fig. 3, Supplemental Fig. 2 & 3, data not shown). As a positive control for *RELA* siRNA efficacy, *RELA* siRNA was sufficient to attenuate SOD2 mRNA and protein levels in IL-1-treated LNCaP, C4-2, and C4-2B cells (Fig. 3, Supplemental Fig. 2) and as a negative control for *RELA* siRNA specificity, *RELA* siRNA did not affect *KEAP1* mRNA levels (Supplemental Fig. 2). Taken together, our data indicates that RELA mediates IL-1 repression of AR in C4-2 or C4-2B cells and to a lesser extent in LNCaP cells.

**Figure 3.**
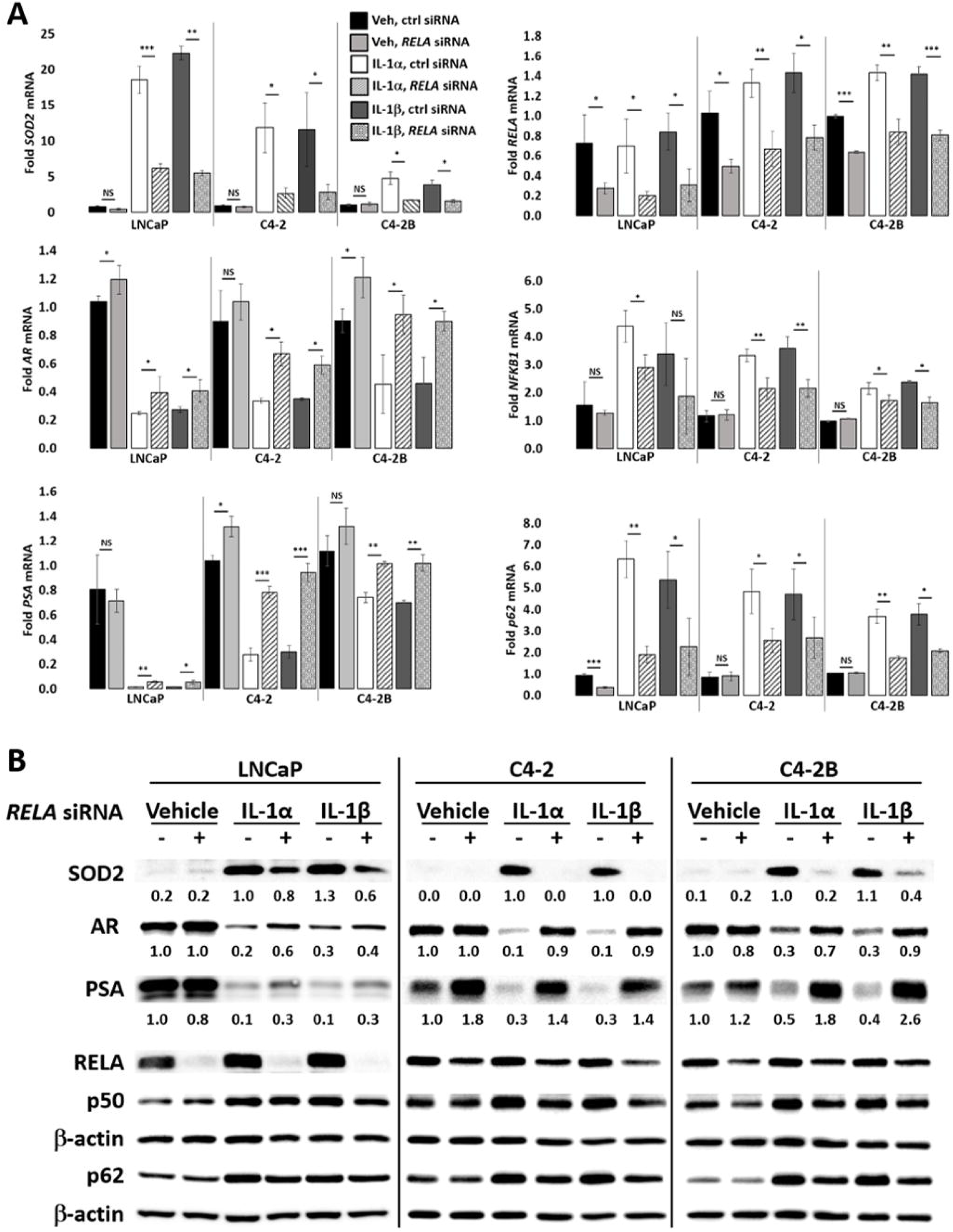
RELA mediates IL-1 repression of AR. (A) RT-QPCR and (B) western blot analysis and densitometry were performed for LNCaP, C4-2, and C4-2B cells transfected with 70 nM of non-targeting or *RELA* siRNA (Dharmacon) for 1 day followed by treatment with vehicle control (Veh), IL-1α, IL-1β for 2 days. LNCaP cells were treated with 0.5 ng/ml IL-1 and C4-2 and C4-2B cells were treated with 25 ng/ml IL-1. *RELA* siRNA attenuates IL-1 downregulation of AR and PSA mRNA and protein and attenuates IL-1 upregulation of SOD2 mRNA and protein. Error bars, ± STDEV of 3 biological replicates; p-value, *≤0.05, **≤0.005, ***≤0.0005. mRNA fold change is normalized to the vehicle control.

**Figure 4.**
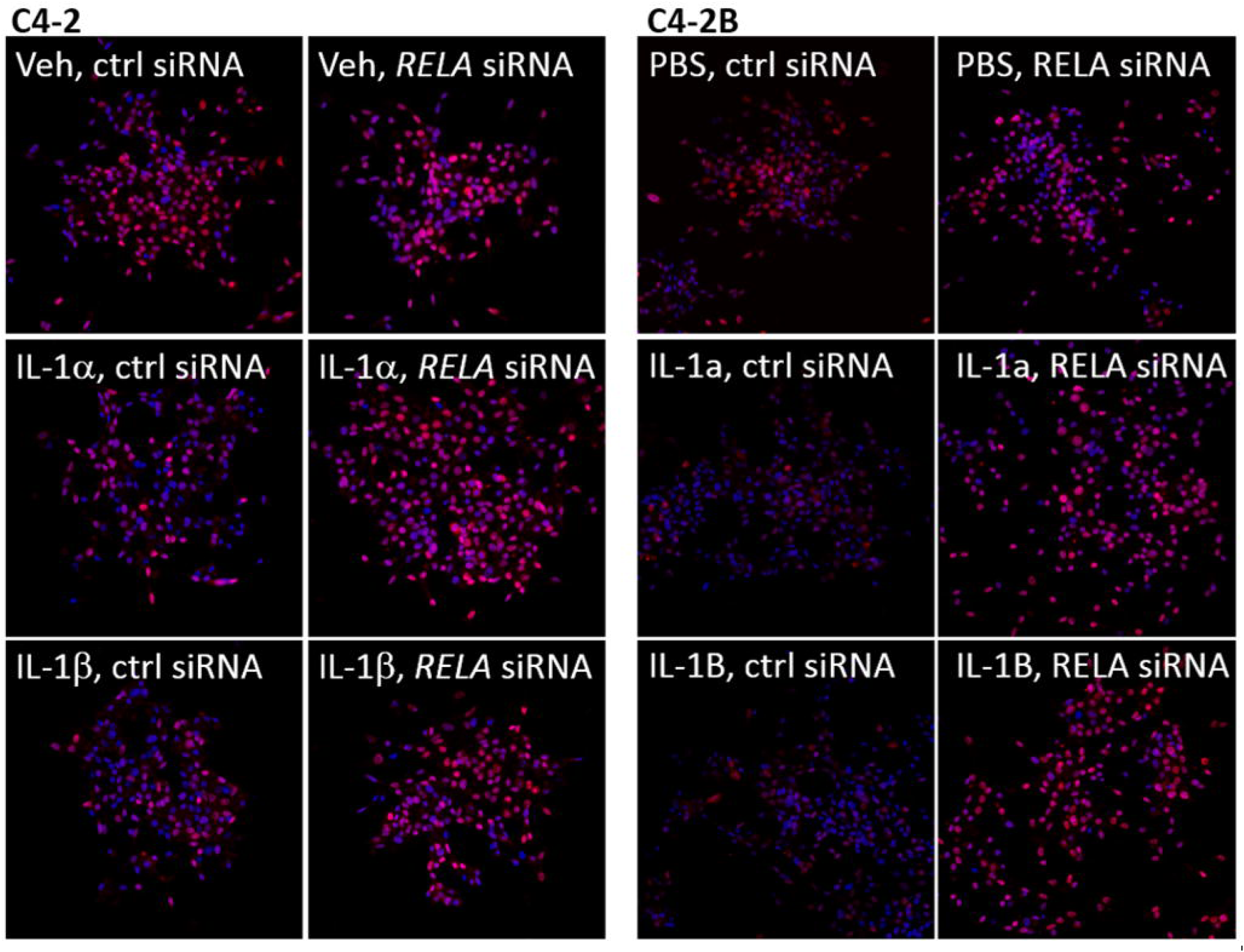
*RELA* siRNA restores nuclear AR accumulation in IL-1-treated PCa cells. C4-2 and C4-2B cells transfected with 70 nM of non-targeting or *RELA* siRNA (Dharmacon) for 1 day followed by treatment with vehicle control (Veh), 25 ng/ml IL-1α, or 25 ng/ml IL-1β for 2 days. Cells were fixed and immunostained for AR (Texas Red) and cell nuclei (DAPI, blue) and imaged at 20X magnification. AR/DAPI merge images are shown for C4-2 (left) and C4-2B (right). RELA siRNA restores AR nuclear accumulation in IL-1-treated cells.

### IL-1 repression of AR levels or activity is specific to the RELA NF-κB family member

Given that the RELA/p50 heterodimer mediates canonical IL-1 signaling^15^ and that *RELA* siRNA downregulates *NFKB1* mRNA and p50 protein in IL-1-treated cells (Fig. 3, Supplemental Fig. 2), we expected that loss of *NFKB1* would also attenuate IL-1 repression of AR levels and activity. *NFKB1* siRNA alone was not sufficient to attenuate IL-1 repression of AR or PSA levels, nor sufficient to block IL-1 induction of SOD2 mRNA or protein levels in LNCaP, C4-2, or C4-2B cells (Fig. 5). Similar results were obtained for *RELB*, *c-REL*, and *NFKB2* siRNA (Supplemental Fig. 4-6). Taken together, our data suggest that RELA is the NF-κB family member that is necessary and sufficient for IL-1-mediated AR repression.

**Figure 5.**
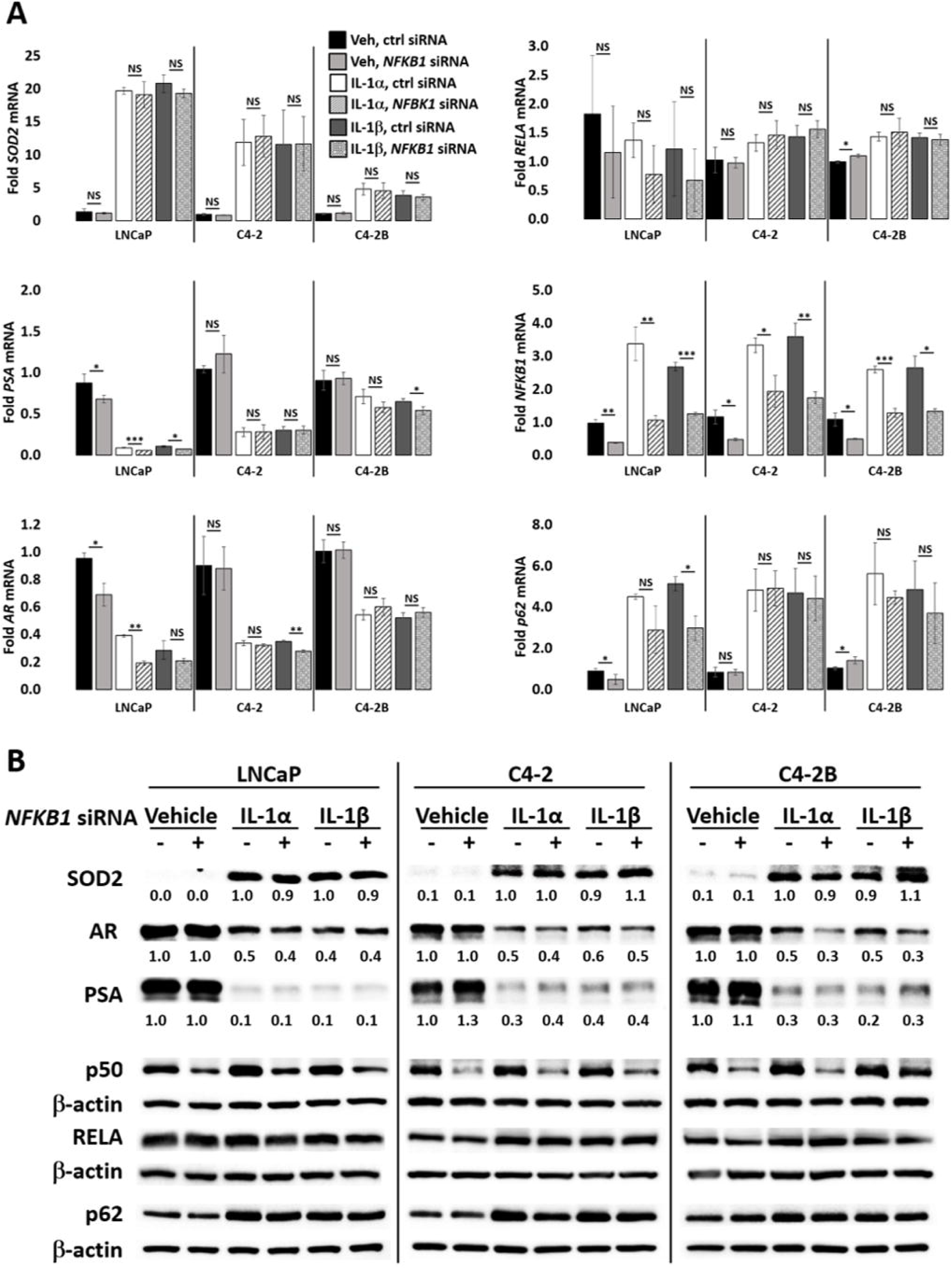
NFKB1(p50) does not mediate IL-1 repression of AR. (A) RT-QPCR and (B) western blot analysis and densitometry were performed for LNCaP, C4-2, and C4-2B cells transfected with 70 nM of non-targeting or *NFKB1* siRNA for 1 day followed by treatment with vehicle control (Veh), IL-1α, IL-1β for 2 days. LNCaP cells were treated with 0.5 ng/ml IL-1 and C4-2 and C4-2B cells were treated with 25 ng/ml IL-1. *NFKB1* siRNA does not attenuate IL-1 downregulation of AR and PSA mRNA and protein, nor does it attenuate IL-1 upregulation of SOD2 mRNA and protein. Error bars, ± STDEV of 3 biological replicates; p-value, *≤0.05, **≤0.005, ***≤0.0005. mRNA fold change is normalized to the vehicle control.

### p62 does not mediate IL-1 repression of AR levels or activity

IL-1 upregulates p62 accumulation in PCa cell lines^6, 14^, *p62* is an IL-1-RELA target gene (Fig. 3), and p62 is a known mediator of IL-1-induced NF-κB activation^16–18^. To investigate if p62 mediates IL-1-RELA repression of AR, IL-1-treated LNCaP, C4-2, or C4-2B cells were transfected with *p62* siRNA. *p62* siRNA was not sufficient to attenuate IL-1 repression of AR or PSA levels or to block IL-1 induction of SOD2 levels in LNCaP, C4-2, or C4-2B cells (Fig. 6). Thus, while IL-1-RELA signaling induces p62 mRNA and protein accumulation, p62 does not appear to be necessary for IL-1-RELA repression of AR levels or activity in LNCaP, C4-2, or C4-2B cells. Interestingly, *p62* siRNA consistently upregulates PSA mRNA and protein in C4-2 and C4-2B cells under basal growth conditions, but *p62* siRNA does not affect PSA or AR levels or AR activity in the presence of IL-1 (Fig. 6). Thus, p62 function includes PSA repression, but independent of IL-1 signaling.

**Figure 6.**
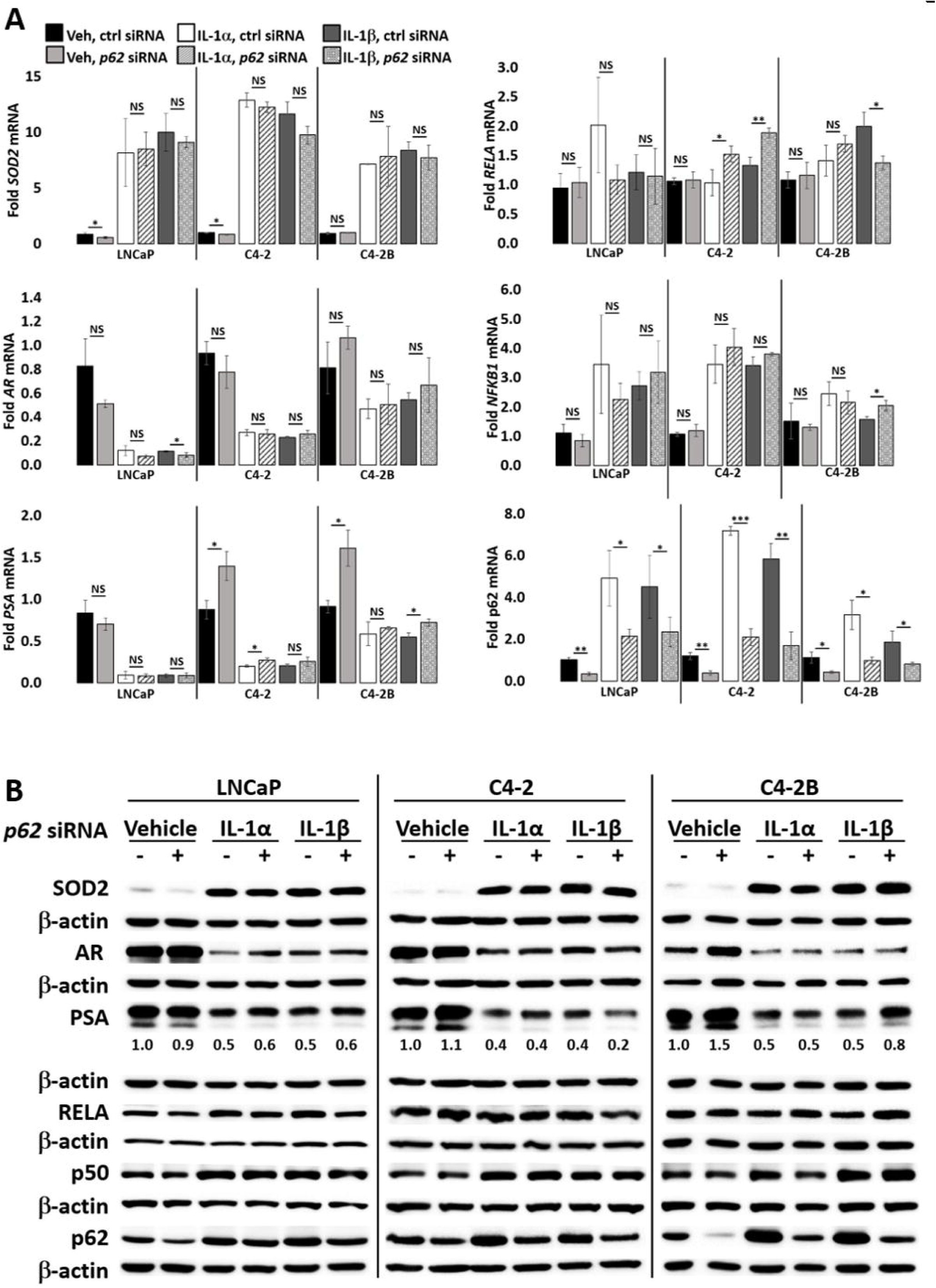
p62 does not mediate IL-1 repression of AR. (A) RT-QPCR and (B) western blot analysis and densitometry were performed for LNCaP, C4-2, and C4-2B cells transfected with 70 nM of non-targeting or *p62* siRNA for 1 day followed by treatment with vehicle control (Veh), IL-1α, IL-1β for 2 days. LNCaP cells were treated with 0.5 ng/ml IL-1 and C4-2 and C4-2B cells were treated with 25 ng/ml IL-1. *p62* siRNA does not attenuate IL-1 downregulation of AR and PSA mRNA and protein, nor does it attenuate IL-1 upregulation of SOD2 mRNA and protein. *p62* siRNA significantly upregulates *PSA* mRNA in C4-2 and C4-2B cells under control growth conditions. Error bars, ± STDEV of 3 biological replicates; p-value, *≤0.05, **≤0.005, ***≤0.0005. mRNA fold change is normalized to the vehicle control.

## DISCUSSION

### IL-1-RELA represses AR mRNA and protein levels

*RELA* overexpression in LNCaP cells has been shown to induce *AR* mRNA accumulation and AR nuclear accumulation^19, 20^; conversely, co-silencing of *RELA* and *p50* reduced AR protein levels in LNCaP cells^19^. Another study found that *p52* overexpression, but not *RELA* overexpression, induces AR nuclear accumulation and activity in LNCaP cells, while neither *p52* nor *RELA* overexpression affected AR mRNA levels^21^. Finally, counter to AR activation by *RELA* or *p52* overexpression, *c-REL* overexpression was shown to block AR activity in LNCaP cells^22^. Thus, NF-κB regulation of AR expression and activity is complex and context-specific.

Notably, the studies described above mimic constitutive NF-κB activity through forced overexpression. We instead, silenced basal or IL-1-induced NF-κB activity in LNCaP, C4-2, or C4-2B cells. We did not observe a change in AR levels or AR activity under basal growth conditions by any NF-κB family member and, using two different pools of oligos, only RELA repressed AR levels and activity and only in the presence of IL-1. RELA is the canonical mediator of IL-1-induced NF-κB signaling^15^, so it is not surprising that RELA would mediate IL-1 repression of AR levels and activity.

Like IL-1, Tumor Necrosis Factor Alpha (TNFα) signals through RELA/p50^15^. TNFα was shown to repress *AR* expression in a NFκB-dependent^23^ or RELA-dependent manner^24^ in LNCaP cells. As we observed for IL-1-treated cells, siRNA loss of *NFKB1* did not affect *AR* mRNA levels in TNFα-treated cells^24^; however, p50 did bind the *AR* promoter in TNFα-treated cells, along with RELA and the HDAC and NCOR co-repressors^24^. It remains to be investigated if IL-1 signaling represses *AR* expression through a signaling mechanism similar to TNFα and/or if RELA indirectly, for example through micro RNAs, downregulates *AR* mRNA levels. Indeed, it has been shown that TNFα reduces *AR* mRNA stability posttranscriptionally^25^. RELA can also form homodimers to regulate transcription^15^; thus, it is also possible that RELA mediates IL-1 repression of AR levels and activity independent of p50.

### IL-1-RELA signaling differs among androgen-dependent LNCaP versus androgen-independent C4-2 and C4-2B cells

We confirmed that the C4-2 and C4-2B cell lines are less dependent on androgen than the LNCaP cell line and we discovered that C4-2 and C4-2B cells are less sensitive to IL-1 signaling than LNCaP cells, yet C4-2 and C4-2B cells are more dependent on RELA-mediated repression of AR levels and activity in response to IL-1 signaling. Interestingly, C4-2 and C4-2B are also less sensitive to TNFα than LNCaP cells and reduced sensitivity is, in part, dependent on reduced basal levels of the TNFα receptor associated protein, TRADD^23^ and we find that C4-2 and C4-2B cells have reduced basal levels of the NFκB p50 transcription factor subunit which likely contributes to reduced IL-1 and TNFα sensitivity. Taken together, our and other published results suggest that during PCa disease progression, cell response to inflammation evolves. To better understand the molecular mechanisms underlying IL-1-RELA signaling in AR-dependent versus AR-independent PCa cells, it will be informative to identify the differences in co-factors, chromatin markers, and chromatin structure at the AR locus in IL-1-treated LNCaP, C4-2, and C4-2B cells.

Our results suggest that changes can occur in IL-1-RELA signaling during PCa disease progression that could promote AR independence and resistance to AR-targeting anti-androgens by selecting for cells that adapt to survive with low or no AR accumulation or activity. Indeed, studies show that inflammation, including IL-1 accumulation, inversely correlate with AR levels or activity in PCa disease progression^7, 8^. Interestingly, while IL-1 has been found to inversely correlate with AR levels or activity, constitutive RELA accumulation appears to positively correlate with biochemical relapse^26, 27^, presumably due to AR activity, in PCa disease progression. In this context, RELA accumulation may reflect evolved constitutive RELA signaling that is independent of cell response to exogenous IL-1.

### *p62* silencing does not prevent IL-1-RELA repression of AR level or activity

Using *p62* siRNA, we were able to reduce *p62* levels to at least the levels observed in the *RELA* knockdowns of IL-1-treated cells and, thus, we would have expected to detect some attenuation of IL-1-induced *AR* repression in *p62*-silenced cells if p62 were required. *p62* silencing did not affect *AR* mRNA levels in IL-1-treated cells. Likewise, the RELA target gene, *SOD2*, did not change in *p62*-silenced IL-1-treated cells, further supporting that RELA can function independently of p62 in response to IL-1. Our results were unexpected given that p62 has been shown to mediate IL-1 activation of NF-κB activity in HepG and HEK293 cells^16–18^; but our results may reflect underlying differences in cell lines and/or treatment conditions.

Interestingly, *p62* silencing did significantly upregulate *PSA* mRNA levels in C4-2 and C4-2B cells under basal growth conditions, suggesting the p62 promotes the repression of basal *PSA* levels and/or basal AR activity. While we did not detect an effect on PSA mRNA or protein levels in LNCaP cells transfected with *p62* siRNA, the sensitivity of RNA sequencing does reveal that *p62* siRNA upregulates *PSA* mRNA levels in LNCaP cells (data not shown). Thus, as observed for IL-1-RELA signaling, LNCaP, C4-2 and C4-2B cells may have differential response to p62 signaling that correlates with castration resistance.

Our *RELA* siRNA data indicate that RELA mediates IL-1 induction of p62 mRNA and protein accumulation. *RELA* siRNA, however, had no effect on p62 levels under basal growth conditions. Thus, p62 repression of basal *PSA* levels and/or basal AR activity is independent of RELA activity. The molecular mechanism and functional significance of p62 repression of *PSA* is unknown but should be explored given that PSA is prognostic for PCa diagnosis and treatment response^28^.

## CONCLUSION

Cumulative evidence suggests that IL-1 promotes androgen and AR independence in PCa. Elevated IL-1 is associated with disease progression^8, 29^, IL-1 accumulation inversely correlates with AR activity in PCa patient tissue^8^, IL-1 signaling down regulates AR levels and activity^3–7^, and chronic IL-1 exposure promotes anti-androgen resistance^5^. Androgen deprivation can induce tumor microenvironment inflammation, including IL-1 accumulation, by promoting immune cell infiltration and PCa cell IL-1 secretion^30, 31^. Thus IL-1 can select for treatment resistant PCa cells, for example following therapy, by selecting for PCa cells that adapt to low or no AR accumulation or activity.

In the era of anti-androgen use, there has been an increase in castration resistant patients with low or no AR activity^32^; the functional or correlative significance of IL-1-RELA signaling has yet to be determined in these patients. Our data shows that IL-1-induced RELA activity can attenuate AR accumulation and activity in PCa cells and, thus, further exploration and clarification of IL-1-RELA signaling in PCa disease progression and treatment resistance is warranted to identify alternative therapeutic targets to AR.

## ACKNOWLEDGEMENTS

For their advice and support throughout this process, we would like to thank all the members of the labs of Drs. Nikki Delk and Chao Xing. We would also like to acknowledge financial support for the Delk lab from the University of Texas at Dallas and financial support from the National Institutes of Health (NIH/NCI R21CA175798 (Delk); NIH/NCI K01CA160602 (Delk); NIH UL1TR001105 (Xing)).

## FIGURE LEGENDS

**Supplemental Table 1. RELA Reactome.** Complete list of molecules in RELA network (Fig. 1) and their relationships with RELA. The relationships are determined from the Ingenuity Knowledge Data Base.

**Supplemental Figure 1.**
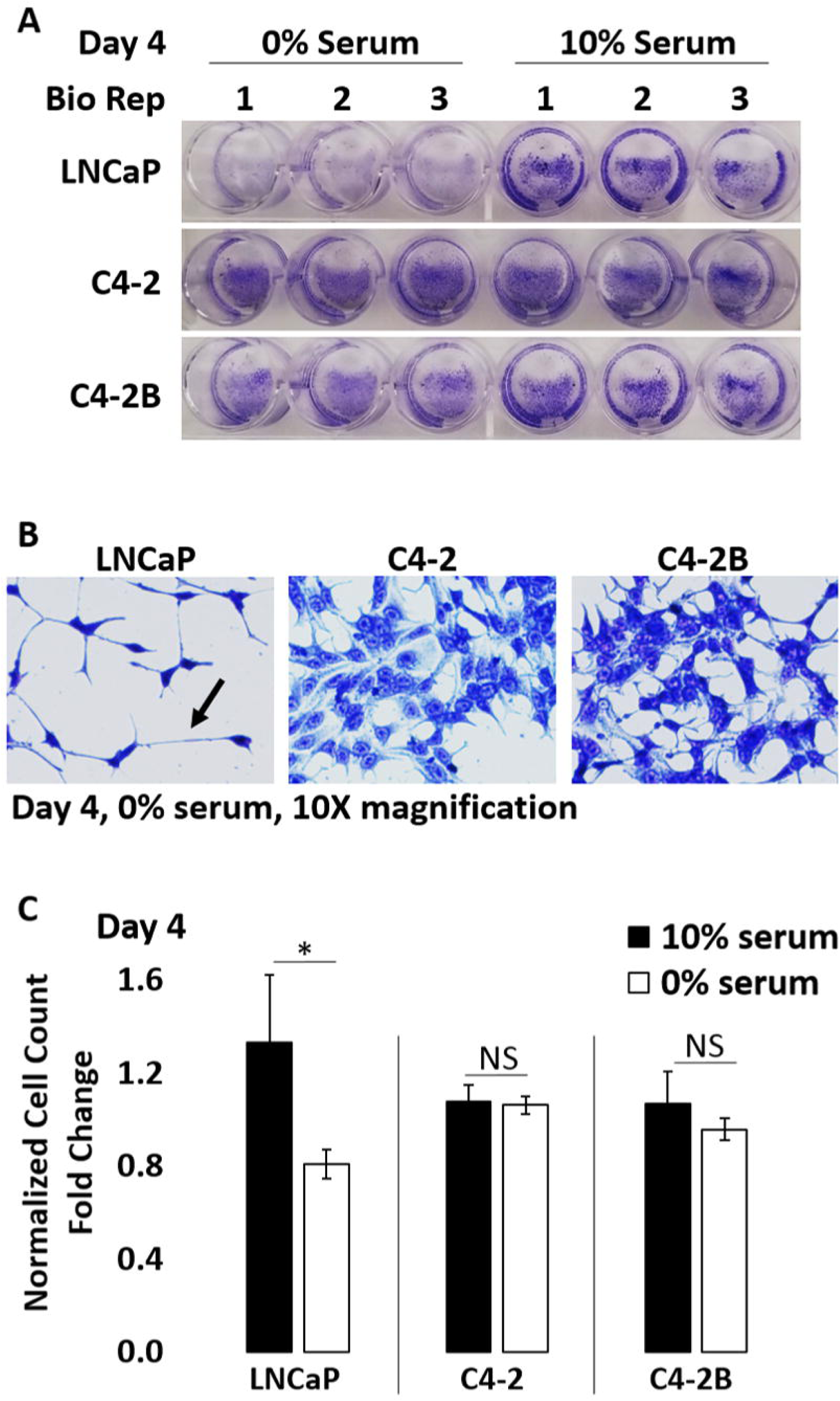
The LNCaP PCa cell line series shows differential sensitivity to serum starvation. LNCaP, C4-2, and C4-2B PCa cells were grown in medium containing 0% or 10% serum for 4 days and (A, B) stained with crystal violet (CV) and imaged or (C) the cells counted and graphed. Serum starvation significantly reduces cell number (A, C) in the LNCaP cells and induces cell process extension (arrow) (B). Thus, LNCaP cells are more sensitive to serum starvation than C4-2 or C4-2B cells. Error bars, ± STDEV of 3 biological replicates; p-value, *≤0.05. Cell counts were normalized to 10% serum treatment control.

**Supplemental Figure 2.**
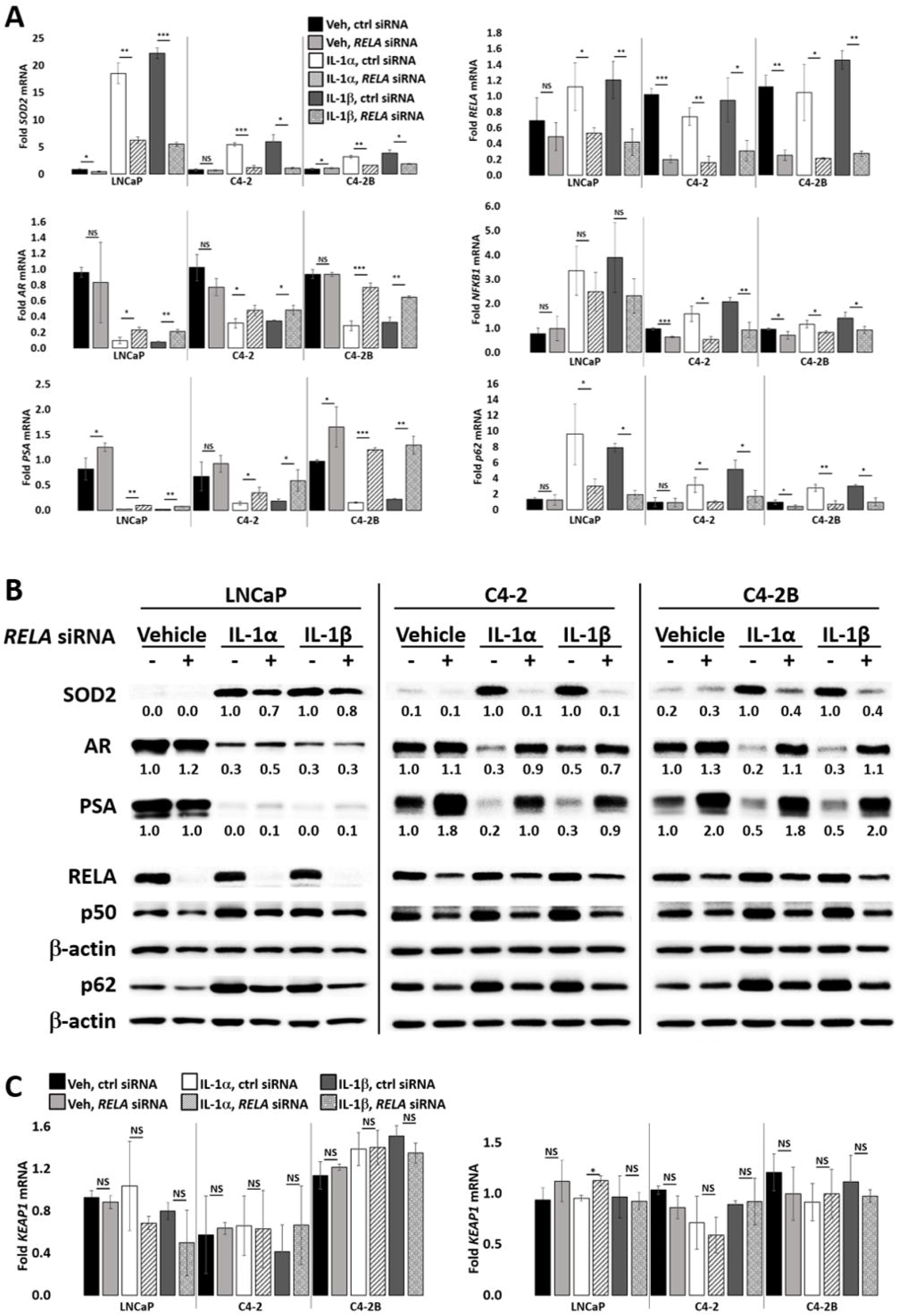
RELA-mediated AR Repression is reproducible and specific. (A) RT-QPCR and (B) western blot analysis and densitometry were performed for LNCaP, C4-2, and C4-2B cells transfected with 70 nM of non-targeting or *RELA* siRNA (Origene) for 1 day followed by treatment with vehicle control (Veh), IL-1α, IL-1β for 2 days. LNCaP cells were treated with 0.5 ng/ml IL-1 and C4-2 and C4-2B cells were treated with 25 ng/ml IL-1. As observed for Dharmacon RELA siRNA (Fig. 3), *RELA* siRNA attenuates IL-1 downregulation of AR and PSA mRNA and protein and attenuates IL-1 upregulation of SOD2 mRNA and protein. (C) LNCaP, C4-2, and C4-2B cells were treated with 25 ng/ml IL-1 and transfected with 70 nM of non-targeting or *RELA* siRNA (Dharmacon, left graph; Origene, right graph) for 1 day followed by treatment with vehicle control (Veh), IL-1α, IL-1β for 2 days and analyzed by RT-QPCR for a non-target gene, *KEAP1*. *RELA* siRNA does not affect *KEAP1* mRNA levels. Error bars, ± STDEV of 3 biological replicates; p-value, *≤0.05, **≤0.005, ***≤0.0005. mRNA fold change is normalized to the vehicle control.

**Supplemental Figure 3.**
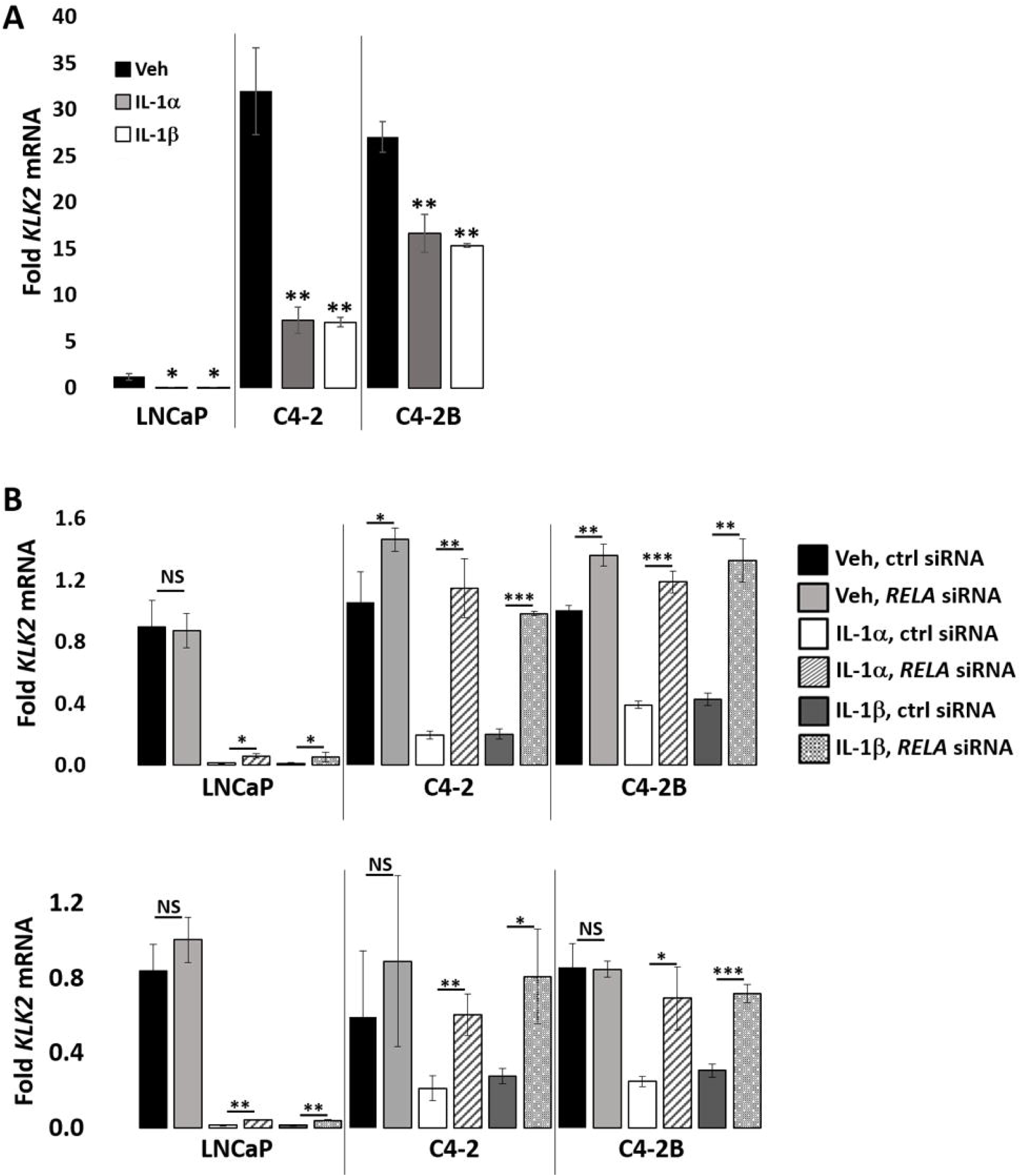
*RELA* siRNA attenuates IL-1 repression of the AR target gene, *KLK2*. (A) RT-QPCR was performed for LNCaP, C4-2, and C4-2B cells treated for 2 days with vehicle control (Veh, V), 25 ng/ml IL-1α, or 25 ng/ml IL-1β and analyzed for the AR target gene, *KLK2*. IL-1 repress *KLK2* mRNA accumulation, which is basally high in C4-2 and C4-2B cells. (B) RT-QPCR was performed for LNCaP, C4-2, and C4-2B cells were transfected with 70 nM of non-targeting or *RELA* siRNA (Dharmacon, top graph; Origene, bottom graph) for 1 day followed by treatment with vehicle control (Veh) or 5 ng/ml IL-1 (LNCaP) or 25 ng/ml IL-1 (C4-2, C4-2B) for 2 days. *RELA* siRNA attenuates IL-1 downregulation of *KLK2* mRNA. Error bars, ± STDEV of 3 biological replicates; p-value, *≤0.05, **≤0.005. mRNA fold change is normalized to LNCaP vehicle control in (A) and normalized to cell line vehicle control in (B).

**Supplemental Figure 4.**
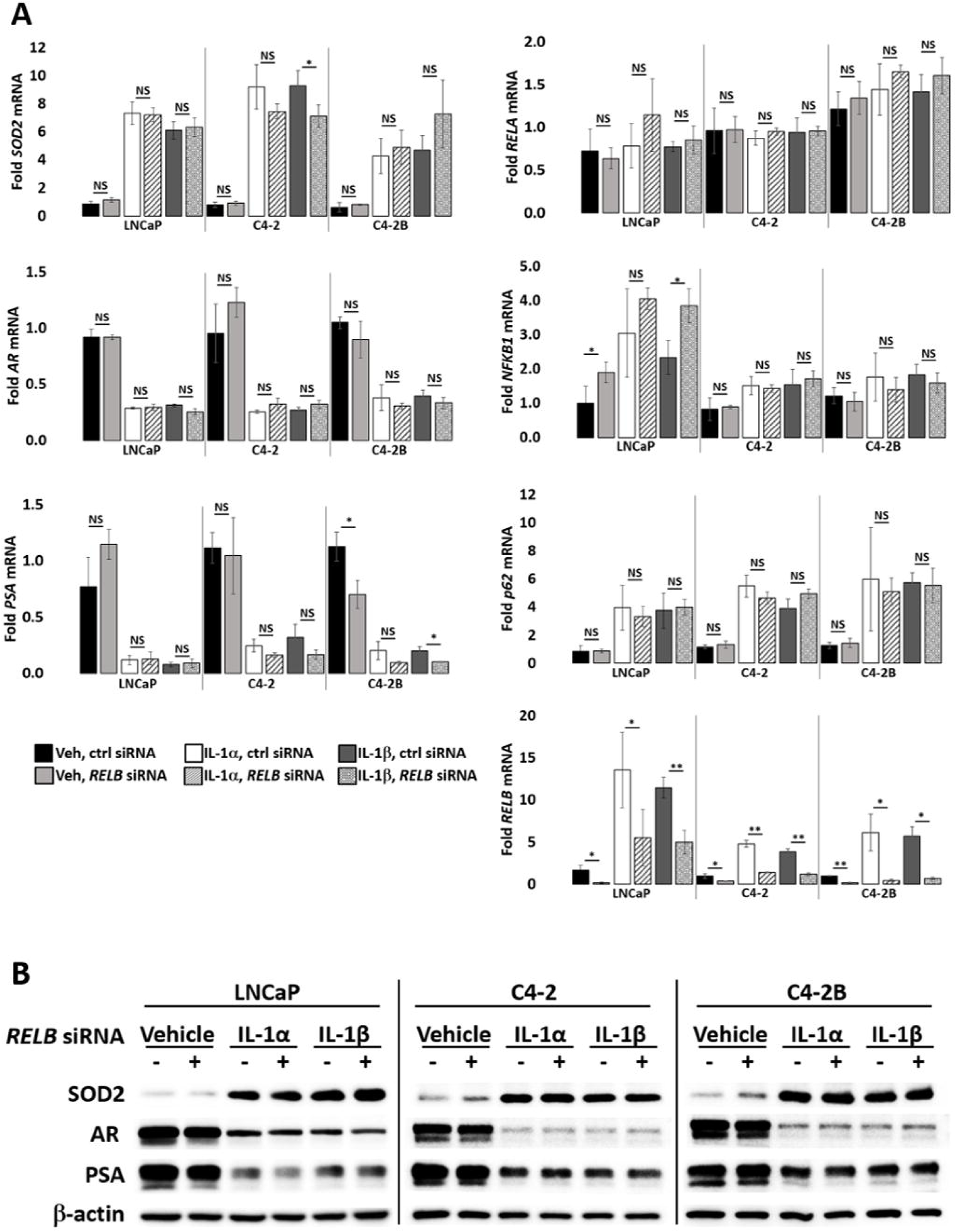
RELB does not mediate IL-1 repression of AR. (A) RT-QPCR and (B) western blot analysis were performed for LNCaP, C4-2, and C4-2B cells transfected with 100 nM of non-targeting or *RELB* siRNA for 1 day followed by treatment with vehicle control (Veh), IL-1α, IL-1β for 2 days. LNCaP cells were treated with 0.5 ng/ml IL-1 and C4-2 and C4-2B cells were treated with 25 ng/ml IL-1. *RELB* siRNA does not attenuate IL-1 downregulation of AR and PSA mRNA and protein, nor does it attenuate IL-1 upregulation of SOD2 mRNA and protein. Error bars, ± STDEV of 3 biological replicates; p-value, *≤0.05, **≤0.005. mRNA fold change is normalized to the vehicle control.

**Supplemental Figure 5.**
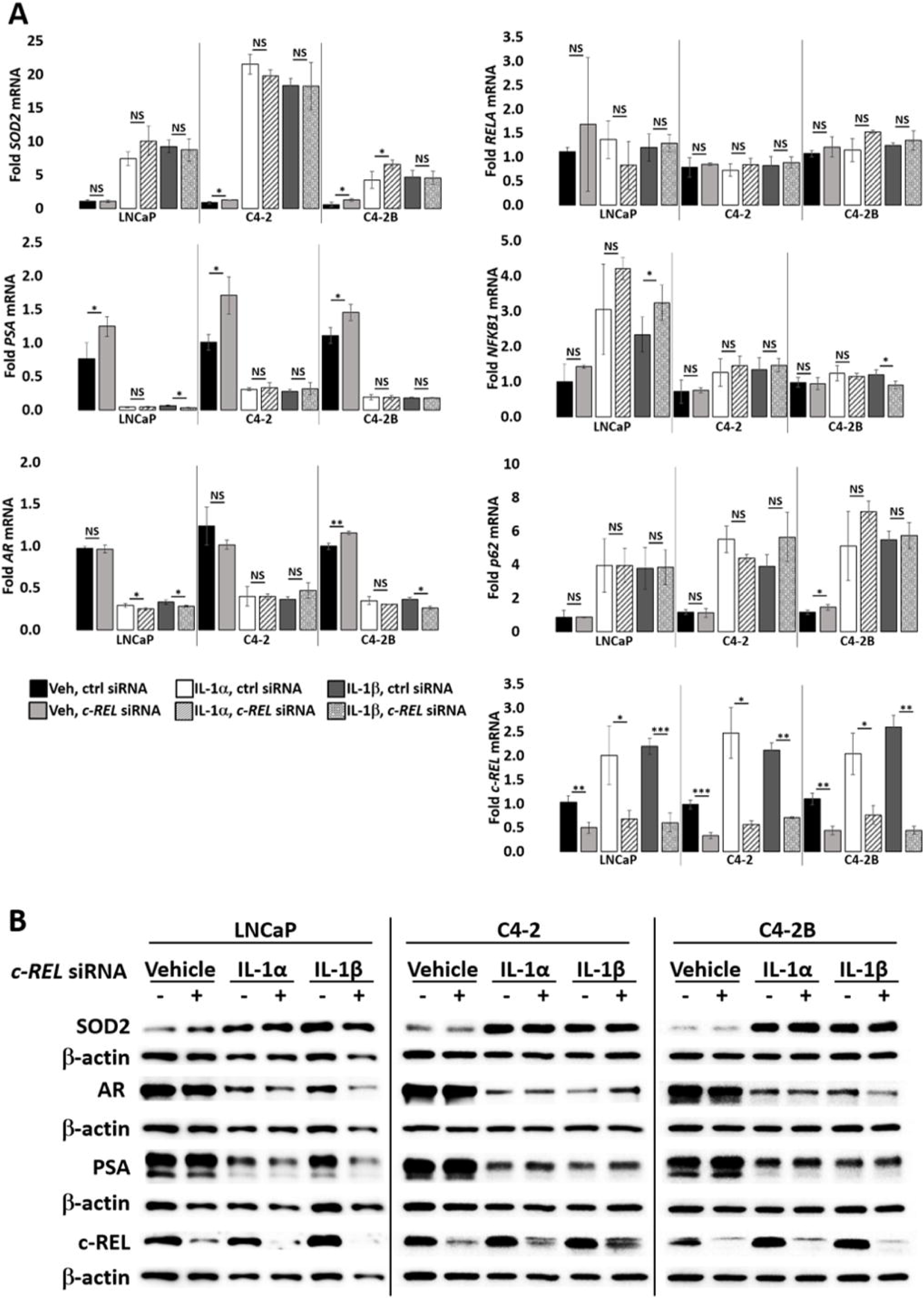
c-REL does not mediate IL-1 repression of AR. (A) RT-QPCR and (B) western blot analysis were performed for LNCaP, C4-2, and C4-2B cells transfected with 100 nM of non-targeting or *c-REL* siRNA for 1 day followed by treatment with vehicle control (Veh), IL-1α, IL-1β for 2 days. LNCaP cells were treated with 0.5 ng/ml IL-1 and C4-2 and C4-2B cells were treated with 25 ng/ml IL-1. *c-REL* siRNA does not attenuate IL-1 downregulation of AR and PSA mRNA and protein, nor does it attenuate IL-1 upregulation of SOD2 mRNA and protein. Error bars, ± STDEV of 3 biological replicates; p-value, *≤0.05, **≤0.005, ***≤0.0005. mRNA fold change is normalized to the vehicle control.

**Supplemental Figure 6.**
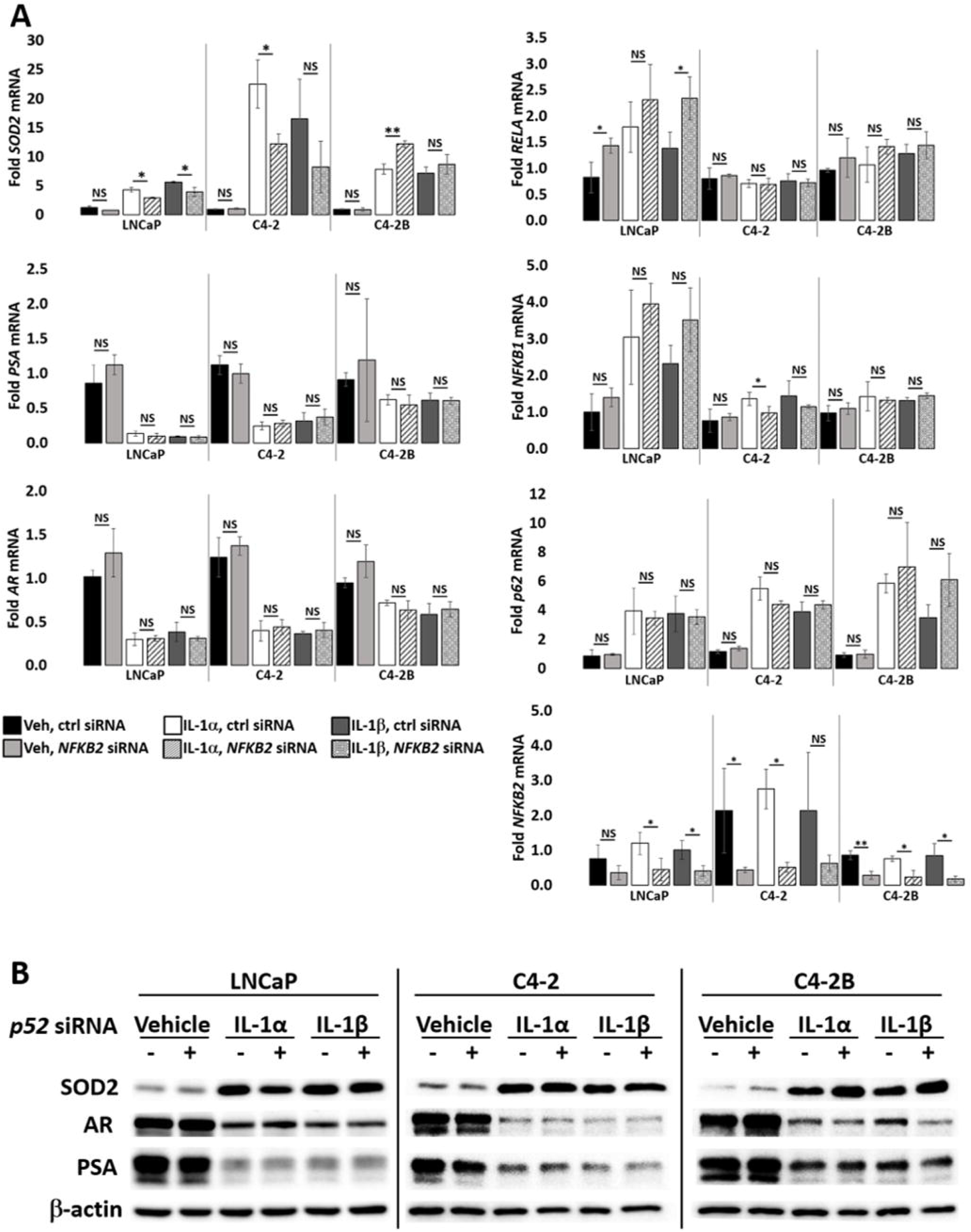
NFKB2/p52 does not mediate IL-1 repression of AR or induction of p62. (A) RT-QPCR and (B) western blot analysis were performed for LNCaP, C4-2, and C4-2B cells transfected with 100 nM of non-targeting or *NFKB2* siRNA for 1 day followed by treatment with vehicle control (Veh), IL-1α, IL-1β for 2 days. LNCaP cells were treated with 0.5 ng/ml IL-1 and C4-2 and C4-2B cells were treated with 25 ng/ml IL-1. *NFKB2* siRNA does not attenuate IL-1 downregulation of AR and PSA mRNA and protein, nor does it attenuate IL-1 upregulation of SOD2 mRNA and protein. Error bars, ± STDEV of 3 biological replicates; p-value, *≤0.05, **≤0.005, ***≤0.0005. mRNA fold change is normalized to the vehicle control.

