## Supplemental Table 1 for "RELA is Sufficient to Mediate Interleukin-1 (IL-1) Repression of Androgen Receptor Expression and Activity in LNCaP Disease Progression Model"

| From Molecule(s) | Relationship Type | To Molecule(s) |
|---|---|---|
| ATF3 | protein-protein interactions | RELA |
| ATF3 | regulation of binding | RELA |
| CUX1 | modification | RELA |
| CUX1 | protein-protein interactions | RELA |
| CYLD | activation | RELA |
| FOXM1 | expression | RELA |
| FOXO3 | activation | RELA |
| FOXO3 | expression | RELA |
| FOXO3 | protein-protein interactions | RELA |
| FOXO3 | translocation | RELA |
| HOXA10 | expression | RELA |
| NFKBIZ | protein-protein interactions | RELA |
| NFKBIZ | regulation of binding | RELA |
| NOTCH3 | activation | RELA |
| PIAS3 | protein-protein interactions | RELA |
| PIAS3 | regulation of binding | RELA |
| PIAS3 | transcription | RELA |
| PML | protein-protein interactions | RELA |
| PML | regulation of binding | RELA |
| POU2F1 | expression | RELA |
| POU2F1 | protein-protein interactions | RELA |
| REL | expression | RELA |
| REL | protein-DNA interactions | RELA |
| REL | protein-protein interactions | RELA |
| RELA | activation | REL |
| RELA | activation | RELA |
| RELA | activation | SMAD3 |
| RELA | expression | ACTA2 |
| RELA | expression | AHR |
| RELA | expression | BACE1 |
| RELA | expression | BACH2 |
| RELA | expression | BCL2 |
| RELA | expression | BCL3 |
| RELA | expression | BEX2 |
| RELA | expression | BIRC3 |
| RELA | expression | BRCA2 |
| RELA | expression | CASP8 |
| RELA | expression | CCL2 |
| RELA | expression | CCL20 |
| RELA | expression | CD274 |
| RELA | expression | CD59 |
| RELA | expression | CDKN2A |
| RELA | expression | CFB |
| RELA | expression | CFTR |
| RELA | expression | CXCL2 |
| RELA | expression | CXCL5 |

| | | |
|---|---|---|
| RELA | expression | CXCL8 |
| RELA | expression | CYP2C9 |
| RELA | expression | DDIT3 |
| RELA | expression | DUSP1 |
| RELA | expression | EGR1 |
| RELA | expression | EPAS1 |
| RELA | expression | ERAP1 |
| RELA | expression | ERAP2 |
| RELA | expression | EZH2 |
| RELA | expression | FOS |
| RELA | expression | GRK5 |
| RELA | expression | GSTA1 |
| RELA | expression | HIF1A |
| RELA | expression | HSP90B1 |
| RELA | expression | ICAM1 |
| RELA | expression | IGF1 |
| RELA | expression | IGF1R |
| RELA | expression | IKBKE |
| RELA | expression | IL1B |
| RELA | expression | IL32 |
| RELA | expression | IRF1 |
| RELA | expression | JUNB |
| RELA | expression | KLF10 |
| RELA | expression | MICA |
| RELA | expression | MMP1 |
| RELA | expression | MST1R |
| RELA | expression | MYC |
| RELA | expression | NFKB1 |
| RELA | expression | NFKB2 |
| RELA | expression | NFKBIA |
| RELA | expression | NFKBIE |
| RELA | expression | NMI |
| RELA | expression | NOX1 |
| RELA | expression | NR4A2 |
| RELA | expression | OAS2 |
| RELA | expression | P2RY2 |
| RELA | expression | PDE4B |
| RELA | expression | PLAU |
| RELA | expression | PLK1 |
| RELA | expression | PPARG |
| RELA | expression | PTGES |
| RELA | expression | PTGS2 |
| RELA | expression | RELA |
| RELA | expression | RELB |
| RELA | expression | SAA1 |
| RELA | expression | SDC4 |
| RELA | expression | SERPINE2 |

| | | |
|---|---|---|
| RELA | expression | SOD2 |
| RELA | expression | TANK |
| RELA | expression | TGM2 |
| RELA | expression | TLR2 |
| RELA | expression | TNF |
| RELA | expression | TNFAIP2 |
| RELA | expression | TNFAIP3 |
| RELA | expression | TNIP1 |
| RELA | expression | TSLP |
| RELA | expression | TWIST1 |
| RELA | expression | UBD |
| RELA | expression | UGT1A1 |
| RELA | expression | VEGFA |
| RELA | expression | WASF3 |
| RELA | localization | STAT3 |
| RELA | modification | RELA |
| RELA | molecular cleavage | FOXO3 |
| RELA | molecular cleavage | RELA |
| RELA | phosphorylation | RELA |
| RELA | phosphorylation | SMAD3 |
| RELA | protein-DNA interactions | ALCAM |
| RELA | protein-DNA interactions | APP |
| RELA | protein-DNA interactions | AR |
| RELA | protein-DNA interactions | B2M |
| RELA | protein-DNA interactions | BACE1 |
| RELA | protein-DNA interactions | BCL10 |
| RELA | protein-DNA interactions | BCL2 |
| RELA | protein-DNA interactions | BCL3 |
| RELA | protein-DNA interactions | BIRC3 |
| RELA | protein-DNA interactions | BIRC5 |
| RELA | protein-DNA interactions | BRCA2 |
| RELA | protein-DNA interactions | C3 |
| RELA | protein-DNA interactions | CCL2 |
| RELA | protein-DNA interactions | CCL20 |
| RELA | protein-DNA interactions | CD44 |
| RELA | protein-DNA interactions | CD59 |
| RELA | protein-DNA interactions | CIITA |
| RELA | protein-DNA interactions | CTGF |
| RELA | protein-DNA interactions | CXCL2 |
| RELA | protein-DNA interactions | CXCL8 |
| RELA | protein-DNA interactions | DUSP1 |
| RELA | protein-DNA interactions | E2F1 |
| RELA | protein-DNA interactions | EGR1 |
| RELA | protein-DNA interactions | ELF3 |
| RELA | protein-DNA interactions | ERAP1 |
| RELA | protein-DNA interactions | ERAP2 |
| RELA | protein-DNA interactions | F3 |

| | | |
|---|---|---|
| RELA | protein-DNA interactions | FN1 |
| RELA | protein-DNA interactions | GRK5 |
| RELA | protein-DNA interactions | HES1 |
| RELA | protein-DNA interactions | ICAM1 |
| RELA | protein-DNA interactions | IER3 |
| RELA | protein-DNA interactions | IFNGR2 |
| RELA | protein-DNA interactions | IGFBP2 |
| RELA | protein-DNA interactions | IL15RA |
| RELA | protein-DNA interactions | IL1B |
| RELA | protein-DNA interactions | IL1RN |
| RELA | protein-DNA interactions | IL7R |
| RELA | protein-DNA interactions | IRF1 |
| RELA | protein-DNA interactions | LTB |
| RELA | protein-DNA interactions | MICA |
| RELA | protein-DNA interactions | MMP1 |
| RELA | protein-DNA interactions | MST1R |
| RELA | protein-DNA interactions | MUC1 |
| RELA | protein-DNA interactions | MYB |
| RELA | protein-DNA interactions | MYC |
| RELA | protein-DNA interactions | NFKB1 |
| RELA | protein-DNA interactions | NFKB2 |
| RELA | protein-DNA interactions | NFKBIA |
| RELA | protein-DNA interactions | NFKBID |
| RELA | protein-DNA interactions | NFKBIE |
| RELA | protein-DNA interactions | NOD2 |
| RELA | protein-DNA interactions | NOX1 |
| RELA | protein-DNA interactions | P2RY2 |
| RELA | protein-DNA interactions | PI3 |
| RELA | protein-DNA interactions | PLAU |
| RELA | protein-DNA interactions | PLK1 |
| RELA | protein-DNA interactions | PRKCD |
| RELA | protein-DNA interactions | PSMB9 |
| RELA | protein-DNA interactions | PTGS2 |
| RELA | protein-DNA interactions | REL |
| RELA | protein-DNA interactions | RELA |
| RELA | protein-DNA interactions | RELB |
| RELA | protein-DNA interactions | SLC1A2 |
| RELA | protein-DNA interactions | SOD2 |
| RELA | protein-DNA interactions | SOX9 |
| RELA | protein-DNA interactions | STAT5A |
| RELA | protein-DNA interactions | TAP1 |
| RELA | protein-DNA interactions | TAPBP |
| RELA | protein-DNA interactions | TGM2 |
| RELA | protein-DNA interactions | TLR2 |
| RELA | protein-DNA interactions | TNF |
| RELA | protein-DNA interactions | TNFAIP2 |
| RELA | protein-DNA interactions | TNFAIP3 |

| | | |
|---|---|---|
| RELA | protein-DNA interactions | TNIP1 |
| RELA | protein-DNA interactions | TSLP |
| RELA | protein-DNA interactions | VASP |
| RELA | protein-DNA interactions | VEGFA |
| RELA | protein-DNA interactions | VHL |
| RELA | protein-DNA interactions | VIM |
| RELA | protein-protein interactions | AFF1 |
| RELA | protein-protein interactions | ATF3 |
| RELA | protein-protein interactions | ATF4 |
| RELA | protein-protein interactions | BCL11B |
| RELA | protein-protein interactions | BRCA1 |
| RELA | protein-protein interactions | CEBPD |
| RELA | protein-protein interactions | CTNNB1 |
| RELA | protein-protein interactions | CUX1 |
| RELA | protein-protein interactions | DAXX |
| RELA | protein-protein interactions | EZH2 |
| RELA | protein-protein interactions | FOXO3 |
| RELA | protein-protein interactions | HES6 |
| RELA | protein-protein interactions | ILF2 |
| RELA | protein-protein interactions | IRF2 |
| RELA | protein-protein interactions | IRF9 |
| RELA | protein-protein interactions | KLF4 |
| RELA | protein-protein interactions | LEF1 |
| RELA | protein-protein interactions | MECOM |
| RELA | protein-protein interactions | NFKBIZ |
| RELA | protein-protein interactions | NPM1 |
| RELA | protein-protein interactions | NRIP1 |
| RELA | protein-protein interactions | PHB2 |
| RELA | protein-protein interactions | PIAS3 |
| RELA | protein-protein interactions | PML |
| RELA | protein-protein interactions | POU2F1 |
| RELA | protein-protein interactions | REL |
| RELA | protein-protein interactions | RELA |
| RELA | protein-protein interactions | SMAD1 |
| RELA | protein-protein interactions | SMAD3 |
| RELA | protein-protein interactions | STAT3 |
| RELA | protein-protein interactions | TAF4B |
| RELA | regulation of binding | ATF4 |
| RELA | regulation of binding | RELA |
| RELA | transcription | APP |
| RELA | transcription | AR |
| RELA | transcription | B2M |
| RELA | transcription | BIRC3 |
| RELA | transcription | BLVRA |
| RELA | transcription | CCL20 |
| RELA | transcription | CDKN2A |
| RELA | transcription | CXCL2 |

| | | |
|---|---|---|
| RELA | transcription | CXCL8 |
| RELA | transcription | DDIT3 |
| RELA | transcription | DUSP1 |
| RELA | transcription | GBP1 |
| RELA | transcription | HLA-B |
| RELA | transcription | ICAM1 |
| RELA | transcription | ID1 |
| RELA | transcription | IER3 |
| RELA | transcription | IL1RN |
| RELA | transcription | KRT15 |
| RELA | transcription | MMP1 |
| RELA | transcription | MYC |
| RELA | transcription | NFATC1 |
| RELA | transcription | NFKB1 |
| RELA | transcription | NFKB2 |
| RELA | transcription | NFKBIA |
| RELA | transcription | NFKBIB |
| RELA | transcription | NOD2 |
| RELA | transcription | P2RY2 |
| RELA | transcription | PLAU |
| RELA | transcription | PRDX6 |
| RELA | transcription | PTGS2 |
| RELA | transcription | RELA |
| RELA | transcription | RELB |
| RELA | transcription | SAA2 |
| RELA | transcription | SDC4 |
| RELA | transcription | SOX9 |
| RELA | transcription | TLR2 |
| RELA | transcription | TNF |
| RELA | transcription | TNFAIP3 |
| RELA | translocation | RELA |
| RELA | ubiquitination | RELA |
| RPL7 | protein-protein interactions | RELA |
| RSF1 | protein-protein interactions | RELA |
| RUNX1 | translocation | RELA |
| RUVBL1 | protein-protein interactions | RELA |
| SMAD3 | protein-protein interactions | RELA |
| SQSTM1 | expression | RELA |
| SQSTM1 | translocation | RELA |
| STAT1 | protein-protein interactions | RELA |
| STAT2 | translocation | RELA |
| STAT3 | localization | RELA |
| STAT3 | protein-protein interactions | RELA |
| STAT3 | translocation | RELA |
| TAF4B | protein-protein interactions | RELA |
| TAF9 | protein-protein interactions | RELA |
| TFAP2A | protein-protein interactions | RELA |

| | | |
|---|---|---|
| TFCP2 | protein-protein interactions | RELA |
| TLE1 | protein-protein interactions | RELA |
| TSC22D3 | activation | RELA |
| TSC22D3 | protein-protein interactions | RELA |
| TSC22D3 | regulation of binding | RELA |
| TSC22D3 | translocation | RELA |
| VDR | protein-protein interactions | RELA |
| YAP1 | protein-protein interactions | RELA |
| YBX1 | protein-protein interactions | RELA |
| ZFP36 | activation | RELA |
| ZFP36 | modification | RELA |
| ZFP36 | protein-protein interactions | RELA |
| ZFP36 | translocation | RELA |
